# Evolutionary rate shifts suggest species-specific adaptation events in HIV-1 and SIV

**DOI:** 10.1101/190769

**Authors:** Maoz Gelbart, Adi Stern

## Abstract

The process of molecular adaptation following a cross-species virus transmission event is currently poorly understood. Here, we identified 137 protein sites that experienced deceleration in their rate of evolution along the HIV-1/SIV phylogeny, likely indicating gain-of-function and consequent adaptation. The majority of such events occurred in parallel to cross-species transmission events and varied between HIV-1 groups, indicating independent adaptation strategies. The evolutionary rate decelerations we found were particularly prominent in accessory proteins that counteract host antiviral restriction factors, suggesting that these factors are a major barrier to viral adaptation to a new host. Surprisingly, we observed that the non-pandemic HIV-1 group O, derived from gorillas, exhibited more rate deceleration events than the pandemic group M, derived from chimpanzees. We suggest that the species barrier is higher when the genetic distance of the hosts increases. Our approach paves the way for subsequent studies on cross-species transfers in other major pathogens.

## Introduction

The Human Immunodeficiency Viruses HIV-1 and HIV-2 are the causative agents of AIDS in humans, infecting millions of people worldwide. Both viruses emerged from a clade of lentiviruses known as the Simian Immunodeficiency Virus (SIV), which naturally infect a variety of non-human primate species. HIV in humans arose from several independent transmission events of primate SIVs that resulted in HIV-1 groups M and N (from SIV infecting chimpanzees, SIVcpz), HIV-1 groups O and P (from SIV infecting gorillas, SIVgor), and HIV-2 groups A through H (from SIV naturally infecting sooty mangabeys, SIVsmm) [1-3]. The gorilla infecting lentivirus, SIVgor, is itself a result of a transmission of SIVcpz to gorillas [4, 5]. Phylogenetic analyses date the most common recent ancestors of HIV groups M and O to the beginning of the 20^th^ century, making it a relatively new human pathogen [6, 7]. Similar analyses of SIVgor date the inception of this virus in the western lowland gorilla population somewhere in the 19^th^ century [4]. SIVcpz itself was found to be a transmission from other primates, dated at roughly 1500 and leading to the two lineages of SIVcpz: SIVcpz*ptt* infecting the chimpanzee subspecies *Pan troglodytes troglodytes* of central Africa, and SIVcpz*pts* infecting the *Pan troglodytes schweinfurthii* chimpanzee subspecies of eastern Africa [5, 8-12].

How viruses are able to cross species barrier is a subject of much interest, since many pandemic human viruses arose from zoonosis events such as the influenza strain of “Spanish flu” H1N1, measles virus (MeV), and SARS coronavirus [13]. Due to genetic differences between the hosts, virus adaptation occurs at multiple levels: at the level of entry to target cells; interaction with the host adaptive immune system; interaction with host antiviral restriction factors; and recruitment of host cellular machinery by the virus [14-16]. Protein adaptations are reflected in the history of genomes and may be manifested in changes in the amino acid substitution rates of the adaptive sites, which are expected to be more conserved in the clade where adaptation happened [17]. This is due to new roles gained by these protein sites that constrain their evolution. A reciprocal phenomenon where amino acids are less conserved in one clade than in other clades is also possible, and may either reflect a loss of function or a gain of function manifested as positive diversifying selection. Identification of such “rate shifting sites” can thus reveal virus adaptation events, and is expected to promote a better understanding of cross-species transmissions in general and in the HIV pandemic in particular in this study.

A method for identification of evolutionary rate changes has been previously established and was used to study intra-subtype HIV adaptation events [18]. To date, the lack of SIVcpz and group O full genomic sequences limited the ability to study adaptation across cross-species transmission events. Here, we utilized the growing availability of diverse and full HIV-1 and SIV genomes to identify many sites in various SIVcpz/SIVgor/HIV-1 clades whose evolutionary rate changed across clades [19]. We demonstrate how cross-species transmissions are correlated with abundant evolutionary rate shifts and how known adaptation events are manifested in different amino acid substitution rates between lineages. Based on the rate shift patterns, we suggest previously unknown adaptation events, and highlight the exceptional amount of evolutionary rate shifts observed in HIV-1 group O, possibly due to a more extreme host species barrier.

## Results

We have previously developed the RASER tool to identify sites that display change in the rate of evolution along a given branch of a phylogenetic tree (see example in Figure 1) [18]. In essence, this tool takes a phylogeny and a multiple sequence alignment and contrasts the rates of evolution of amino acids along all branches in the phylogeny, to determine if the evolutionary rates of some amino acids in some branches are better explained by a model that allows for evolutionary rate changes. We queried the Los Alamos HIV database [19] for all available HIV-1, SIVcpz and SIVgor full sequences, from which we took the sequences of all nine HIV-1 proteins. Due to high representation of group M sequences, we downsampled this group to the size of available group O sequences, retaining the strains with the most internal variation (total number of sequences N=126, see Methods). A different downsample of group M sequences to a bigger size was also conducted, to validate results robustness to cohort size variation (N=223, see Methods). All site coordinates are reported based on the HIV-1 subtype B reference strain HXB2 (Methods).

**Figure 1.**
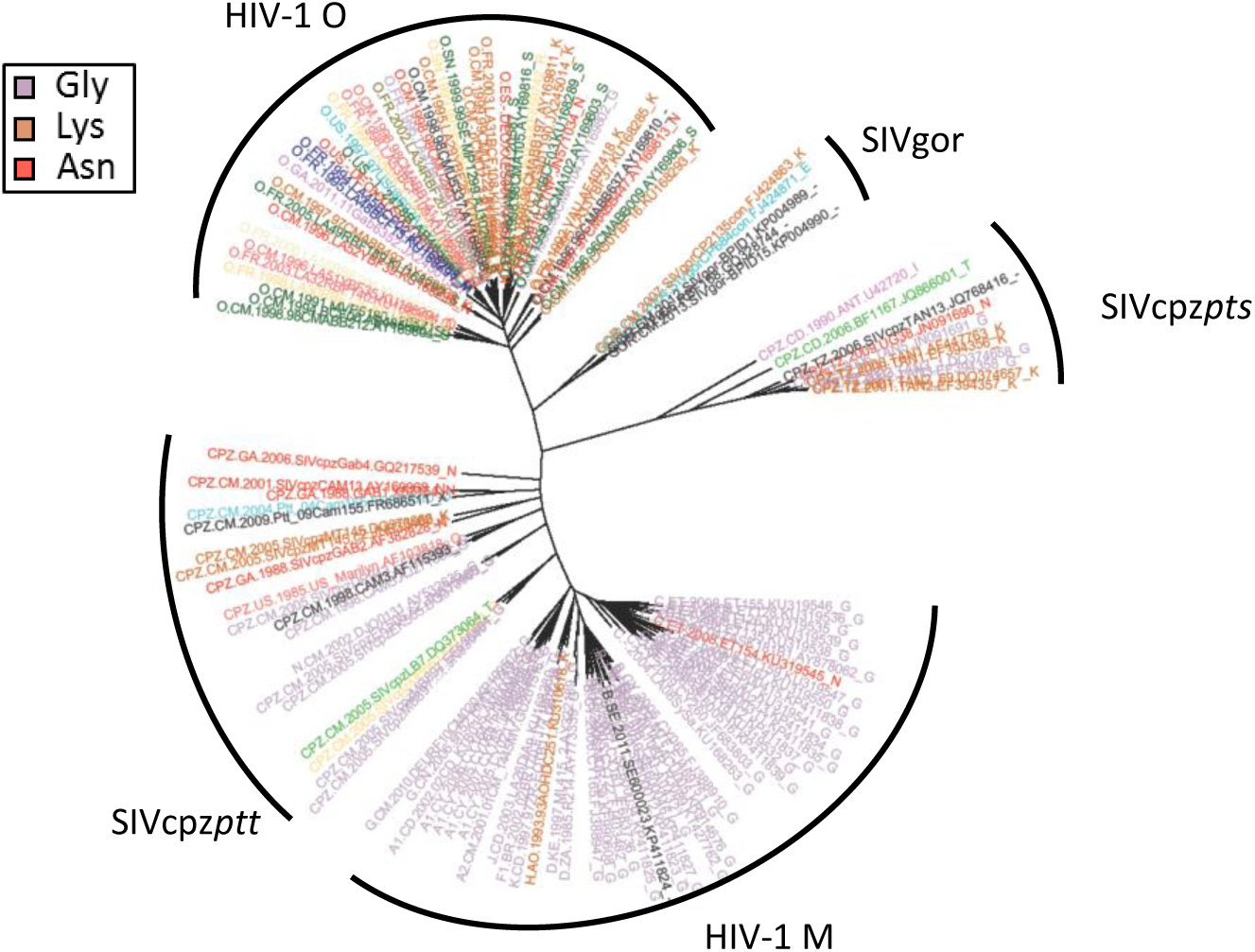
Projection of Env_458_ on the HIV-1/SIV phylogeny. Each leaf (corresponding to an HIV/SIV strain) is color coded based on the amino acid present at Env_458_. The evolutionary rate of Env_458_ was found to be slower in HIV-1 group M than in the rest of the phylogeny.

### Most rate shift events were identified in speciation branches

We first sought to characterize the branches where rate shift events were suggested. A total of 271 rate shifting positions were identified along the phylogeny. 137 of them were identified as rate-deceleration events and an additional 134 were identified as rate-accelerations. As listed in Table 1, 230 out of the 271 rate shifting positions were attributed to four branches: the branch separating group M from SIVcpz, the branch separating group O from SIVgor, the branch separating SIVgor and group O from SIVcpz and the branch separating SIVcpz*pts* from other viruses. Likelihood ratio tests were found to be highly significant in favor of a rate shift model across all HIV proteins (Supplementary Table 2).

**Table 1.** Rate shifts as *percentage from total protein size* for prominent branches, for rate decelerating sites (upper) and rate accelerating sites (lower), colored by intensity. The raw data underlying this table are provided in Supplementary File 2.

| DEC | Structural proteins |  |  | Regulatory proteins |  | Accessory proteins |  |  |  | Total number of mutations |
| --- | --- | --- | --- | --- | --- | --- | --- | --- | --- | --- |
| Branch/protein | Env | Gag | Pol | Rev | Tat | Nef | Vif | Vpr | Vpu |  |
| **Group M** | 1% | 1% | 1% | 1% | 0% | 0% | 0% | 0% | 5% | 28 |
| **Group O** | 2% | 1% | 0% | 8% | 0% | 1% | 1% | 1% | 4% | 46 |
| **SIVgor+P+O** | 0% | 0% | 0% | 0% | 0% | 2% | 1% | 0% | 0% | 14 |
| **SIVcpzpts** | 1% | 1% | 0% | 0% | 0% | 3% | 2% | 1% | 5% | 27 |
| ACC |  |  |  |  |  |  |  |  |  | Total number of mutations |
| Branch/protein | Env | Gag | Pol | Rev | Tat | Nef | Vif | Vpr | Vpu |  |
| **Group M** | 1% | 1% | 0% | 0% | 1% | 1% | 1% | 1% | 1% | 23 |
| **Group O** | 3% | 2% | 1% | 3% | 0% | 1% | 2% | 0% | 1% | 51 |
| **SIVgor+P+O** | 0% | 0% | 0% | 0% | 0% | 2% | 0% | 1% | 1% | 10 |
| **SIVcpzpts** | 1% | 1% | 0% | 0% | 0% | 3% | 1% | 2% | 5% | 31 |

### Different rate deceleration patterns identified in different branches

A special focus was given to rate deceleration events, as they likely indicate a gain of function in a protein. In Figure 2 we mapped the percent of rate shifts in each protein observed in each lineage, corresponding to different virus speciation events. While rate deceleration distribution patterns differed between the different lineages (see Figure 2 and Supplementary Table 1), one common theme was that all prominent branches underwent a significant portion of their rate deceleration events in the Env protein. The next two common rate decelerating proteins were Gag and Pol identified in all four highlighted branches, which is expected given that together with Env these are the longest proteins in HIV/SIV. When controlling for protein size (as in Table 1), a different pattern emerged: Vpu seems to have undergone a disproportionate portion of rate deceleration events, in particular in the branch leading to group M. In general, the rate deceleration patterns for the non-structural proteins seemed to be branch-specific: In the branch leading to SIVgor and group O, Nef and Vif experienced the most relative rate deceleration events. In the branch leading to group M, Vpu was responsible for the most rate deceleration events; and in the branch leading to group O, Rev contributed a significant portion of all rate decelerations for that branch (Supplementary Table 1). The branch separating SIVcpz*pts* from other viruses displayed a more diverse pattern with relatively less rate decelerations in the Env protein and more rate decelerations in the Nef, Vif and Vpu proteins.

**Figure 2.**
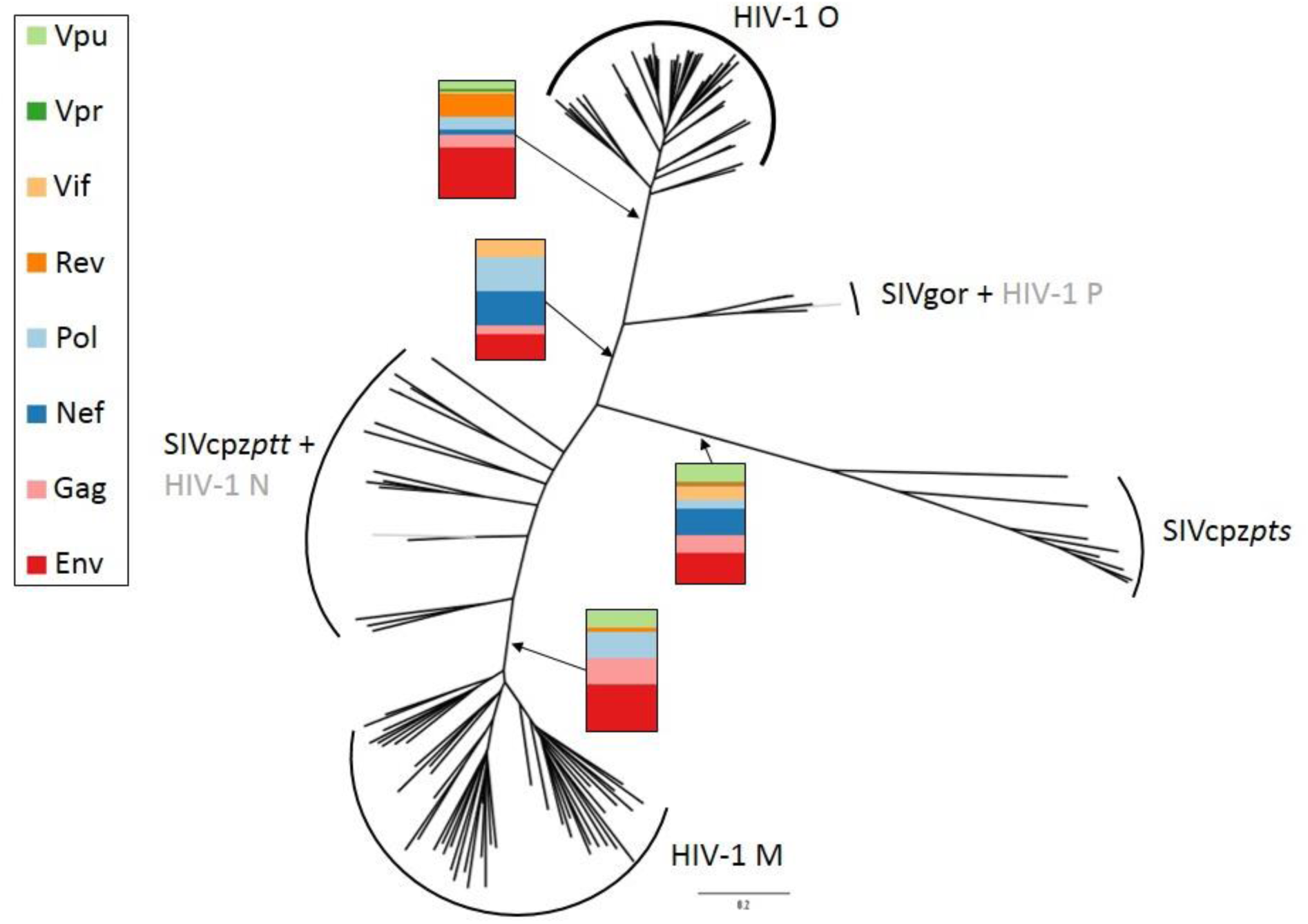
Proposed rate deceleration patterns in prominent branches, shown on the phylogeny. Rate decelerations are shown as percentage from total rate decelerations for each branch.

### Parallel rate shift events in HIV-1 groups M and O

Rate shift events common to several HIV-1 groups are of special interest, since they may better reflect the species barrier in general and not lineage-specific adaptation event. As is evident from Table 2, most parallel rate shift events of HIV-1 groups O and M (or their closest precursor branch) were rate deceleration events, half of which occurred in the highly conserved Pol protein. Notably, the consensus amino acid identity in those parallel-occurring rate deceleration events varied between the M and O clade for all identified positions, suggesting once again that different adaptation strategies occurred in each lineage. Only a single parallel rate acceleration event was identified, in Nef_8_ of HIV-1 groups M and O. This may indicate a relaxation of prior constraint as the SIV allele was maintained in many HIV-1 sequences.

**Table 2.** Rate shifting sites identified independently in parallel in HIV-1 groups M and O or their immediate ancestor.

| Protein | Rate Shifting site | Rate shift type | Branches | Consensus Amino Acids |
| --- | --- | --- | --- | --- |
| Pol | 349 | Deceleration | M, Pre O | M: E, Pre O: P |
| Pol | 406 | Deceleration | Pre M, O | Pre M: S, O: V |
| Pol | 926 | Deceleration | M, Pre O | M: K, Pre O: T |
| Env | 61 | Deceleration | M, O | M: Y, O: T |
| Env | 772 | Deceleration | M, O | M: R, O: S |
| Rev | 71 | Deceleration | Pre M, O | Pre M: V, O: N |
| Nef | 8 | Acceleration | M, O | NA |

NA- Not applicable

### Rate shift events are often parallel to known macromolecular adaptation events

Intense research into the origins of the HIV-1 pandemic and its sources identified several major adaptation events of HIV-1 to its human host. These events include adapting to cellular variations such as differences in host receptor CD4; overcoming host innate immune restriction factors such as Tetherin (BST-2) and APOBEC3G; and of course, evasion from the adaptive immune system [16, 20-25]. We therefore sought to evaluate whether the proposed rate shifts agree with the known adaptation events.

#### Anti Tetherin/BST-2 adaptations

Tetherin/BST-2 is a trans-membrane antiviral restriction factor that disrupts the budding phase of the retroviral lifecycle, thus preventing the infection from spreading [26, 27]. SIVcpz and many more SIV’s antagonize Tetherin through interactions of their nef protein with the cytoplasmic tail of Tetherin [28, 29]. Tetherin is a fast evolving protein, and differs between chimpanzee, gorilla and human [30, 31]. In humans, a significant deletion in the cytoplasmic tail of Tetherin rendered it invulnerable to the SIV’s Nef-based counteraction [29]. Therefore, in order to regain infectivity HIV-1 developed an alternative way to antagonize Tetherin. In groups M and N, Vpu protein adapted to antagonize Tetherin through its transmembrane domain [32]. In group O the virus adapted its Nef protein to counteract the human Tetherin, albeit with less efficiency than in group M [33, 34]. Notably, a single O strain that uses its Vpu protein to encounter human Tetherin has been described [35]. No anti-Tetherin adaptation event is known to have occurred in the rare group P [36]. In gorilla, the Nef protein adapted to encounter its host’s Tetherin but also maintained its ability to encounter the chimpanzee tetherin [37, 38].

Our results support the described anti-Tetherin adaptation events: a dramatically high proportion (14% of all identified group M rate decelerations) were found in the Vpu protein, most of them in the Tetherin-binding domain (Supplementary Table 1 and Figure 3). In addition, in HIV-1 group M the Nef protein was identified with three rate accelerations, supporting the loss of function of its anti-Tetherin activity. This is also supported by a reintroduction experiment of HIV-1 group M strain into chimpanzee in which M-Nef readapted to chimpanzee host, as the mutations required for M-Nef to regain anti-Tetherin activity in chimpanzees (Nef_163_ and Nef_169_) [39] were located proximally to Nef_157_, one of the rate accelerating sites of M-Nef identified in our study. Numerous O-Nef rate acceleration events were found as well, supporting the loss of chimpanzee/gorilla anti-Tetherin function. A preponderance of acceleration and deceleration events were also found in the lineages leading to the ancestor of SIVgor and O and to SIVcpz*pts*; one of those rate deceleration events is at Nef_177_, which is proximal to the C-loop region which has been shown to be related to O-Nef anti-Tetherin activity [40]. Thus, our analysis pinpoints the sites that are likely responsible for the dramatic changes in function of Vpu and Nef throughout SIVcpz and HIV evolution.

**Figure 3.**
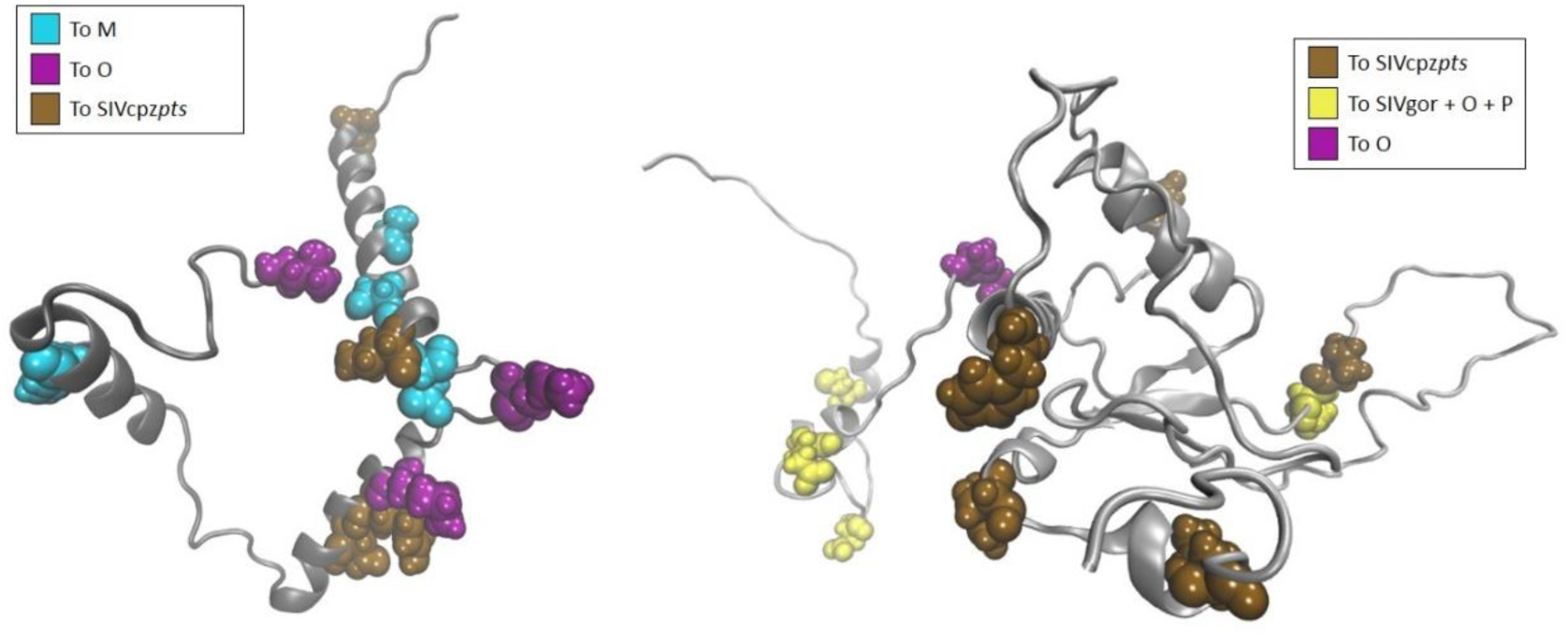
Projection of the identified rate decelerations onto the tertiary structure of Vpu protein (left, PDB ID: 2N28) and Nef protein (right, obtained from [41]) in the major lineages where it was identified.

#### Anti APOBEC3G adaptations

APOBEC3G (A3G) is a broad-range antiviral protein that is packaged into HIV-1 virions, and upon reverse transcription adds C⟶U mutations to the synthesized DNA strand, thus potentially creating nonviable genomes [42]. Lentiviruses use their Vif protein to overcome this factor by degrading it as well as other antiviral proteins from the APOBEC3 family (such as A3F, A3D) [43]. A3G variants of human, chimpanzee and gorilla have been studied and the Vif recognition domain has been identified at A3G residues 126-132 [44]. Of those species, gorilla has a different residue at A3G position 129 as compared to that of human and chimpanzee A3Gs and indeed the SIVgor adaptation event to the different host A3G has been demonstrated by [45, 46]. It has been shown that SIVgor can replicate in the presence of human-, gorilla- and chimpanzee-A3G, but HIV-1 group M or SIVcpz cannot replicate in the presence of gorilla-A3G, indicative of a gain-of-function in the adaptation process of SIVgor to gorilla [46].

In line with these findings, our analysis shows that the branch leading to SIVgor and HIV-1 groups O and P experienced 14% of its rate decelerations in the Vif protein, particularly in positions Vif_73_ and Vif_167_. In HIV-1, Vif_73_ is in close proximity to the A3G-binding motif Vif_69-72_ that is also responsible in-part for interaction with APOBEC3F (which in itself contains differences in the Vif interacting loop between chimpanzee and human/gorilla) [47-49]. O-Vif experienced a single rate deceleration in position Vif_127_, which is located in the Cullin-5 interacting domain [48]. Lack of rate deceleration events in M-Vif suggest no adaptation of this protein in this strain to the human host; this is supported by the observation that SIVcpz-Vif can encounter human A3G [46].

#### Env adaptations

Another major barrier for host jumps lies at the cell entry level. CD4 is the target receptor of SIV and HIV, while CCR5 or CXCR4 act as co-receptors, recognized by the viral Env protein which is exposed to the virus external surface. Previous studies mapped the genetic diversity between different chimpanzees’ CD4 and revealed that there are several differences between chimpanzee subspecies as well as differences between chimpanzee versus gorilla and human, especially in the regions that are in close contact with Env subunit gp120 [50, 51]. Some of those changes affect the glycosylation patterns of CD4, making these differences even more distinguishable [50, 51]. It is therefore likely that HIV-1 had to adapt to recognize these differences in CD4.

Our analysis overall supports a high level of deceleration and acceleration events found in Env, possibly reflecting the changes that occurred in the host CD4. For instance, many of the changes in M-Env occurred in CD4 interacting residues of Env_374_, Env_455_, Env_456_ and Env_458_ [52]. However, we cannot rule out that the very high rate of evolution in Env, driven by the adaptive immune system [53], have led to changes that “mimic” a rate shift. Indeed when accounting for the size of the Env protein, it seems that this protein is less pronounced in its rate shift distribution (Table 1).

#### Gag_30_ human adaptation marker

Position 30 of Gag polyprotein has been previously suggested to be an adaptation event of HIV-1 to its human host, as it diversified from Methionine in SIVcpz*ptt* and SIVgor strains into Lysine or Arginine in many HIV-1 strains [54]. Reintroduction of HIV-1 to chimpanzees resulted in reversion of this position back to the conserved Met of SIVcpz*ptt* and therefore it was suggested as an adaptation event to the human host [54]. Notably, subtype C of group M is conserved for Methionine at Gag_30_, suggesting that adaptation occurred only in some subtypes.

Our results indeed suggest an evolutionary rate shift event in Gag_30_ yet are not conclusive about the branches in which the rate shift happened. This is likely since this site displays a “content shift” rather than a strong “rate shift” (elaborated in the discussion).

**Table 3.** Summary of HIV rate deceleration events related to known antiviral activity.

| Viral Lineage | Viral Protein | Rate decelerating positions identified | Rate deceleration(s) related to the following restriction factor | Function of the restriction factor |
| --- | --- | --- | --- | --- |
| Group M | Vpu | 18, 23, 28, 66 | Tetherin | Prevents virus release from infected cells |
| SIVgor and Group O | Nef | 20, 28, 38, 49, 50, 177 | Tetherin | Prevents virus release from infected cells |
| SIVgor and group O | Vif | 73, 127, 167 | APOBEC3G | Induces hypermutation of viral genomes |

## Discussion

Understanding the molecular changes in a pathogen when adapting to infect a new host species is of high importance. To the best of our knowledge, our study represents the first large-scale approach to detect adaptation events in the transition from non-human primates to humans, and relies on a robust phylogenetic modeling approach. We discovered many lineage-specific adaptation-like events in many proteins of HIV-1, SIVgor and SIVcpz*pts*, and have reported sites in where groups M and O suggested to undergone parallel rate shifts. The majority of sites undergoing rate deceleration were found in the Env protein; when correcting for protein size, the relatively highest number of rate decelerating were found in proteins that adapted to host restriction factors. Accordingly, we suggest that the major common barrier for a host species jump is composed of both the entry stage and the stage where the virus must overcome the first barrier of cellular defenses. Our results further support a differential model of adaptation: groups M and O underwent adaptation in different proteins, and when adaptation occurred at similar sites, the amino-acid was different in both groups. This model is supported by the fact that activity of group M and O proteins are indeed different (e.g., Vpu), and by the fact that each group originated from an SIV from a different primate.

Since group M is the pandemic strain of HIV-1, we initially expected that it would experience the highest number of rate shift events among all HIV-1 groups, indicating more efficient adaptation to its human host. Surprisingly, our analysis revealed that group O had almost twice the amount of rate shift events, despite being a non-pandemic strain that remained localized mainly to infections in west-central Africa [55, 56]. We suggest that the number of rate shift events is not the determinant of a pandemic strain; it may rather reflect the relative height of the species barrier a strain had to overcome, as chimpanzee from which group M originated is genetically closer to human than gorilla from which group O originated [57, 58].

We further noted a large number of rate shifts that occurred in the lineage separating the two subspecies of SIVcpz (SIVcpz*ptt* and SIVcpz*pts*). Presumably these events correspond to adaptations of the virus to the two chimpanzee subspecies, which are genetically divergent. Further research will be required to understand if and how this has affected the adaptation of the virus to the human host.

### Limitations

This study has several limitations. First, the availability of full coverage of SIVcpz and the SIVgor genomes is still low, reducing the statistical power of the analysis. Indeed, we noted that increasing the sample size of group M sequences that are much more available, led to detection of more rate shifting sites (Supplementary File 1). We expect that the availability of additional SIVgor sequences will increase the number of rate shift events that are unique in that clade, possibly revealing more important sites for this lineage. Second, the method that we utilized to identify rate shifts is calibrated to identify dramatic changes in the evolutionary rate along a lineage. Accordingly, it cannot detect sites where a “content shift” occurred, i.e. the amino acid changed and remained conserved in two complementary lineages, since this entails only a minor change in evolutionary rate. This is partially demonstrated in Gag_30_, where the rate of evolution changed mildly, while the content of this site changed between chimpanzee and human viruses.

### Implications

The compiled list of the positions suggested as rate decelerating can now serve as a guide for future functional studies that aim to understand the differences among HIV-1 and SIV proteins. Furthermore, the ability to track adaptation events by utilizing sequence data highlights the power of the method when studying emerging pandemics, strengthening the need to sequence full genomes of pathogens broadly.

### Conclusions

Genetic sequences of viruses and specifically HIV-1 and SIV viruses can be harnessed to identify adaptation events of emerging pathogens to their new host species. Our results suggest that innate immunity serves as a strong barrier for cross-species transmission events, and that this barrier imposed a strong selective pressure for viruses to adapt as they crossed these barriers with increasing efficiency.

## Methods

In order to collect sequences for this study, the Los Alamos HIV sequence database (available online at http://www.hiv.lanl.gov [19]) was queried for HIV-1 sequences from the same strain that spanned all nine HIV-1 open reading frames (ORFs: *gag, pol, vif, vpr, tat, rev, vpu, env* and *nef*) and for SIVcpz and SIVgor strains that spanned the corresponding ORFs (with the exception of vpx). This led to 2004 sequences of HIV-1 group M, 45 sequences of HIV-1 group O, 9 sequences of HIV-1 group N, 2 sequences of HIV-1 group P, 4 sequences of SIVgor and 29 sequences of SIVcpz. Two different datasets were constructed: (i) with a very large number of group M sequences compared to the amount of group O sequences, and (ii) with an equal number of group M and group O sequences. Due to computational reasons, dataset (i) included 200 HIV-1 sequences, most of them group M. In both datasets, the sequences were sampled so that the n most distant strains (in terms of genetic distance) were sampled. Due to extremely high similarity, HIV-1 groups N and P sequences were reduced to a single representative strain from each. The IIIB_LAI strain was added manually as a reference sequence. The results in this study are reported mainly with datasets (ii), chosen since it allows comparing the result from group M and group O. Additional sites found with datasets (i) are reported in Supplementary File 1.

Initial multiple sequence alignments of the nine proteins were performed using PRANK and iteratively improved until convergence [59]. In order to reconstruct of the phylogenetic relationship between the sequences, we concatenated the alignments of Gag, Pol, Vif, Vpr, Tat and Env and provided this as input for PhyML [60]. We next used the reconstructed phylogeny as a guide tree to realign each protein with PRANK. JpHMM was used to validate that the strains used in the analysis are not inter-group recombinants [61].

In order to identify evolutionary rate shifts, we used RASER [18] to analyze each of the nine proteins separately, with the proteome-based phylogeny as input. RASER is a likelihood-based phylogenetic method for detecting a change in site-specific evolutionary rates. First, a likelihood ratio test against a null model of no rate shifts is performed in order to assess if a model enabling rate shifts better fits the data. Next, the posterior probability of rate-shift is calculated at each site and sites with a probability higher than 0.6 are considered here as significant. Finally, for each such site, the method lists the lineages where the rate shift occurred with the highest probability, and further categorizes each sites as undergoing either a rate-deceleration or a rate-acceleration.

In order to test for sequence sampling effects on the identified rate shift patterns, we repeated the analysis with increased amount of group M sequences (n=183), reduced amount of group O sequences (n=13) and no SIVgor sequences, denoted as dataset (i). Analysis revealed more rate decelerations in the branch leading to group M than in dataset (ii) analysis (48 compared to 28, Supplementary File 1). 54% of the positions identified as rate-decelerating in group M in dataset (ii) (n=15) were also identified in dataset (i). Chi-squared tests for differences in group M rate shift distributions showed no significant difference (p=0.11 and 0.87 for rate decelerations and accelerations, respectively), indicating that the patterns of rate shifts between the large group M sample and the smaller group M sample are similar.

## Acknowledgements

The authors would like to thank Dr. Osnat Penn, Prof. Eran Bacharach (Tel Aviv University) and Prof. Tal Pupko (Tel Aviv University) for their contribution to the inception of this study, Prof. Matthias Geyer (University of Bonn) for providing the Nef protein model structure, Dr. Tzachi Hagai and Stern lab members for helpful comments and discussion.

## Competing Interests

The authors declare that they have no competing interests involved with this study.

## Supplementary Figures and Tables

**Supplementary Figure 1.**
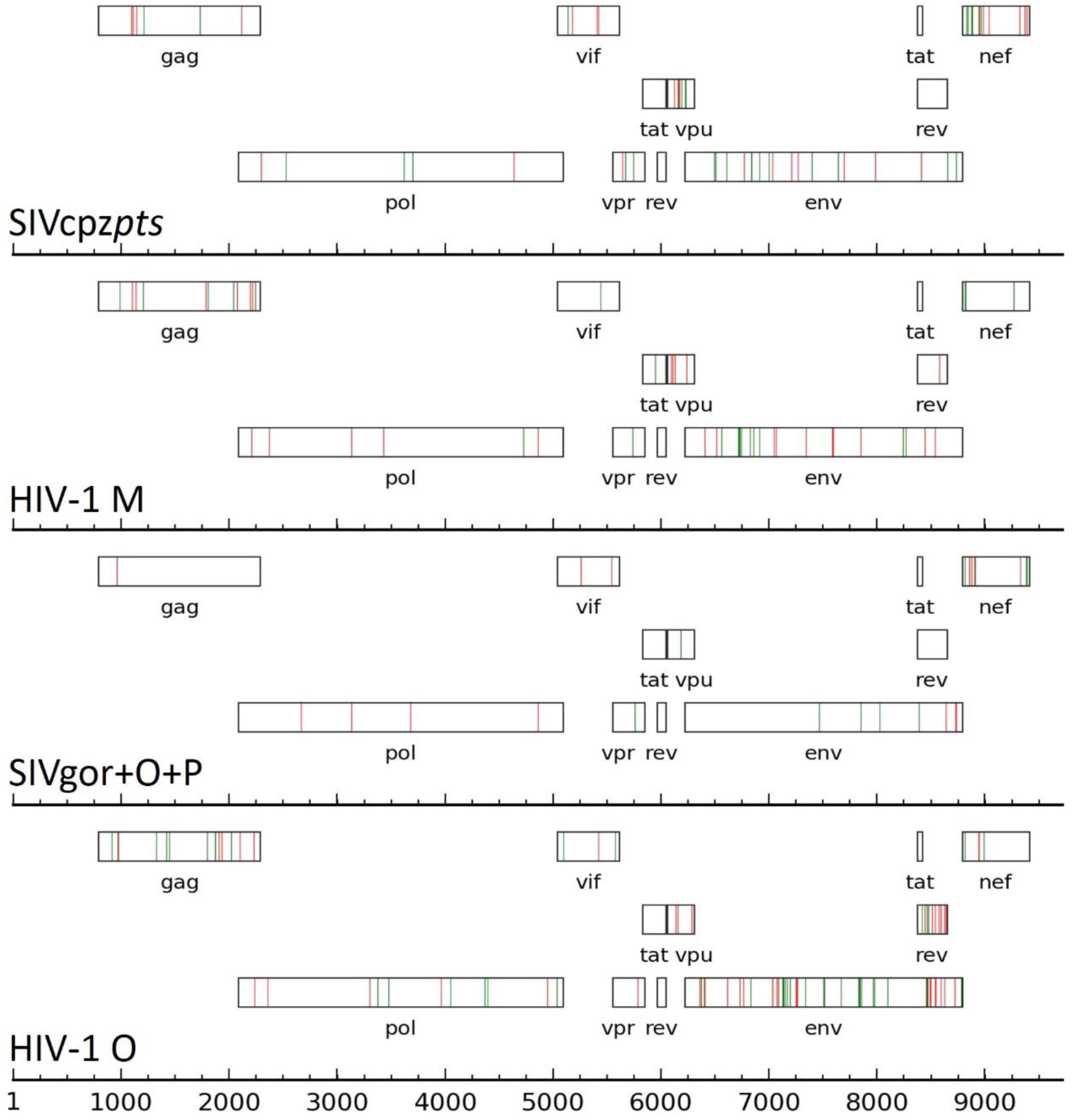
Proposed rate deceleration patterns in prominent branches shown along the genome. Data shown in HXB2 coordinates.

**Supplementary Figure 2.**
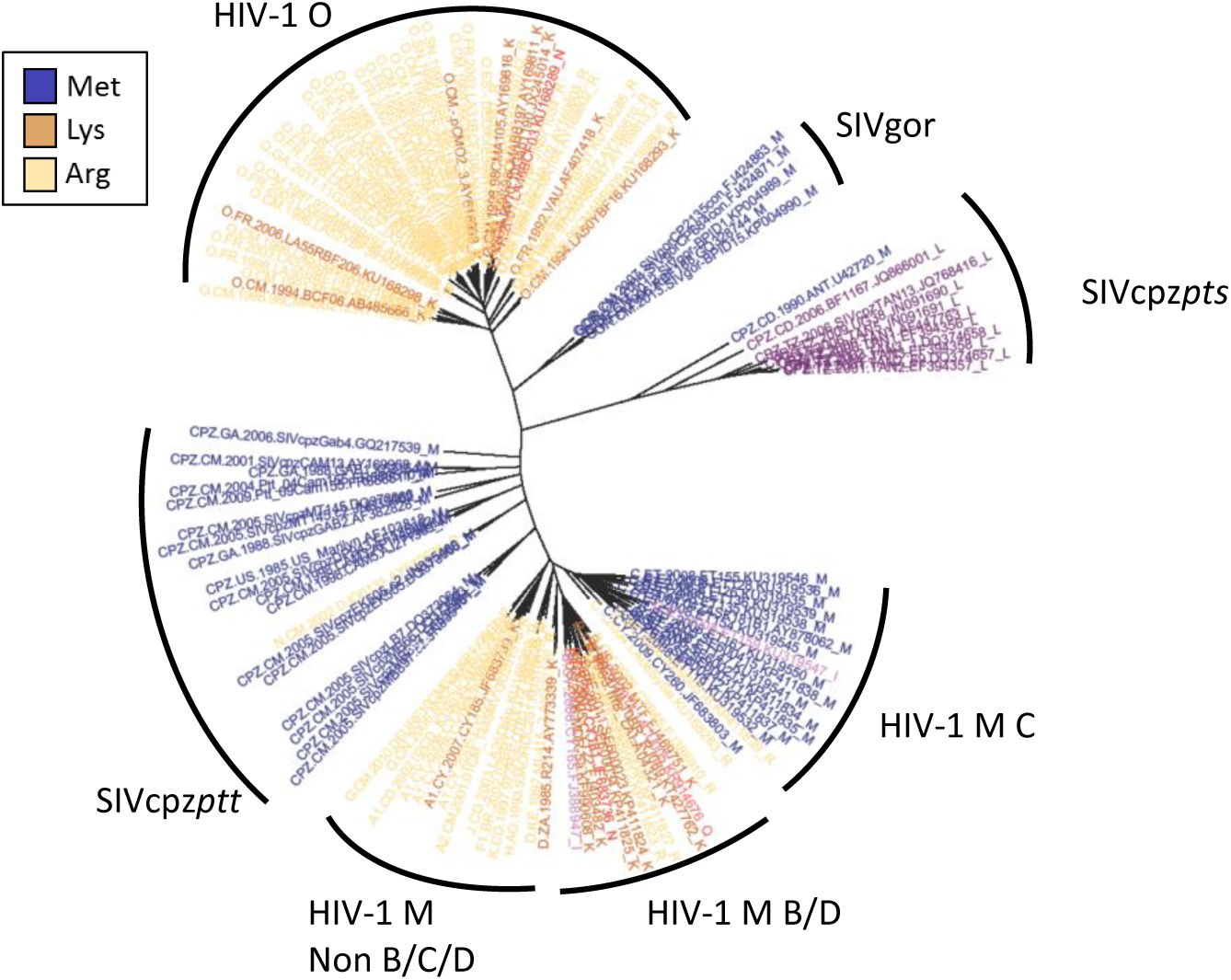
Residues distribution for Gag_30_ along the HIV-1/SIV phylogeny. This position exhibits a pattern more similar to “content shift” than to “rate shift”.

**Supplementary Table 1.**
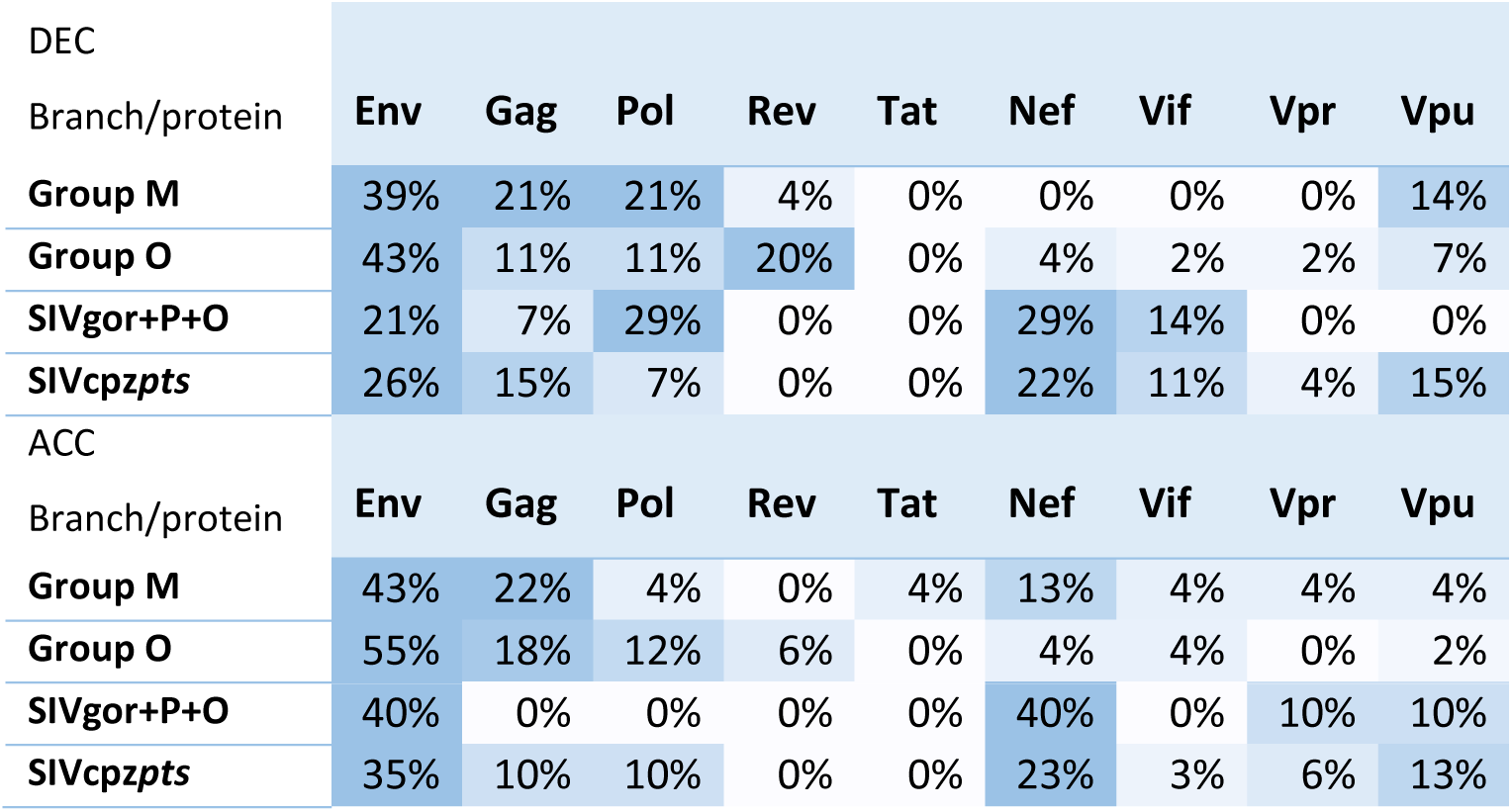
Rate shifts as *percentage from total rate shifts for prominent branches*, for rate decelerating sites (upper) and rate accelerating sites (lower), colored by intensity.

**Supplementary Table 2.** Maximum log-likelihood (LL) values for the analysis of the nine HIV-1/SIVcpz/SIVgor proteins under the rate shift and null models.

| Protein | Rate Shift Model LL | Null Model LL | 2ΔLL | P-value ($\chi^2_3$) |
| --- | --- | --- | --- | --- |
| Gag | -29,478.3 | -29,671.4 | 386 | <10<sup>-50</sup> |
| Pol | -40,813.1 | -41,097.2 | 568 | <10<sup>-100</sup> |
| Vif | -12,677.6 | -12,756.3 | 157 | <10<sup>-30</sup> |
| Vpr | -5,328.1 | -5,355.76 | 55 | <10<sup>-10</sup> |
| Tat | -8,918.23 | -8,947.61 | 59 | <10<sup>-10</sup> |
| Rev | -12,662 | -12,733.6 | 143 | <10<sup>-30</sup> |
| Vpu | -10,548.7 | -10,664.1 | 231 | <10<sup>-50</sup> |
| Env | -92,347.6 | -93,028.8 | 1362 | <10<sup>-100</sup> |
| Nef | -15,611.4 | -15,804.1 | 385 | <10<sup>-50</sup> |

## Supplementary and Source Files

### Supplementary File 1.

**HIV/SIV sites identified as rate shifting for dataset (i)**. Positions are provided in HXB2 reference sequence coordinates. Branch number field refer to the branch number outputted by RASER [18]; for ease of reading we provide branch labels for most branches.

### Supplementary File 2.

**HIV/SIV sites identified as rate shifting for dataset (ii)**. Positions are provided in HXB2 reference sequence coordinates. Branch number field refer to the branch number outputted by RASER [18]; for ease of reading we provide branch labels for most branches.

